# The nematode (*Ascaris suum*) intestine is a location of synergistic anthelmintic effects of Cry5B and levamisole

**DOI:** 10.1101/2023.11.20.567786

**Authors:** Paul D. E. Williams, Matthew T. Brewer, Raffi Aroian, Alan P. Robertson, Richard J. Martin

**Author notes:** Corresponding author: Richard J. Martin, *Email.

## Abstract

A novel group of biocidal compounds are the Crystal 3D (Cry) and Cytolytic (Cyt) proteins produced by *Bacillus thuringiensis* (Bt). Some Bt Cry proteins have a selective nematocidal activity, with Cry5B being the most studied. Cry5B kills nematode parasites by binding selectively to membrane glycosphingolipids, then forming pores in the cell membranes of the intestine leading to damage. Cry5B selectively targets multiple species of nematodes from different clades and has no effect against mammalian hosts. Levamisole is a cholinomimetic anthelmintic that acts by selectively opening L-subtype nicotinic acetylcholine receptor ion-channels (L-AChRs) that have been found on muscles of nematodes. A synergistic nematocidal interaction between levamisole and Cry5B has been described previously, but the location, mechanism and time-course of this synergism is not known. In this study we follow the timeline of the effects of levamisole and Cry5B on the Ca^2+^ levels in enterocyte cells from the intestine of *Ascaris suum* using fluorescence imaging. The peak Ca^2+^ responses to levamisole were observed after approximately 10 minutes while the peak responses to activated Cry5B were observed after approximately 80 minutes. When levamisole and Cry5B were applied simultaneously, we observed that the responses to Cry5B were bigger and occurred sooner than when it was applied by itself. It is proposed that there is an irreversible cytoplasmic Ca^2+^ overload that leads to necrotic cell-death in the enterocyte that is induced by levamisole opening Ca^2+^ permeable L-subtype nAChRs and the development of Ca^2+^ permeable Cry5B toxin pores in enterocyte plasma membranes. The effects of levamisole potentiate and speed the actions of Cry5B.

## Introduction

Infections by soil-transmitted helminths (STHs), including: the roundworm *Ascaris lumbricoides,* the whipworm *Trichuris trichiura,* and hookworms (*Ancylostoma ceylanicum* and *Necator americanus*) are major medical and public health concerns in many developing countries. It is estimated that 807 million to 1.2 billion people are infected with the large roundworm *Ascaris lumbricoides* [1]. Although rarely fatal, parasitic infections have a detrimental effect on morbidity, reducing worker productivity by 6.3 million **D**isability **A**djusted **L**ife **Y**ears (DALYs) per year and have a severe impact on children due to stunting of physical growth, cognitive impairment, malnutrition and anemia. In livestock, infestations lead to reduced food yields, impacting of economic returns and the exacerbation of poverty [2].

Current treatments rely on the use of chemotherapeutics, usually using one of the three major classes of anthelmintics: benzimidazoles (*e.g.,* albendazole/mebendazole), macrocyclic lactones (*e.g.,* ivermectin), or nicotinic cholinergics (*e.g.,* levamisole) because there are no effective vaccines. Resistance to these anthelmintics is frequently observed in domestic animals [3] and there is a real concern that resistance will present in parasites of humans. With the limited number of anthelmintic drugs and the concerns about the development of resistance it is important to: 1) seek novel classes of anthelmintics that have different modes of action and; 2) to identify mechanism-based rational anthelmintic combinations that are more potent and effective.

*Bacillus thuringiensis* (Bt) three-domain crystal (Cry) proteins are a potential novel class of anthelmintic. It is known that Bt Cry proteins form pores in the intestine of insect pests and are used successfully for insect control. Once ingested by insects, these toxins are solubilized in their midgut, and activated by proteases to yield the active toxin. The toxin then binds to various membrane receptors including glycopeptides [4, 5]. The toxin is then transported into the plasma membrane of intestinal cells of the insect where it forms oligomeric pores that cause ion leakage and cell lysis [4]. A major advantage of the Cry proteins is that they are very specific, targeting only invertebrates and are therefore safe for use in vertebrates even at high concentrations. Vertebrates lack these glycoprotein receptors [6–8]. The potency and biosafety of the Bt Cry protein toxins have resulted in them being used widely for insect pest control in transgenic food crops.

A selection of Bt Cry proteins has been identified as being effective at targeting free-living and parasitic nematodes [9–14]. Cry5B is the most studied of these nematode specific Cry proteins. It acts as a pore forming toxin by binding to glycosphingolipid receptors created by BRE-5 [15, 16] and/or cadherin CDH-8 receptors [17]. After binding to the cadherin receptors, the toxin oligomerizes to form the active pore(?) [17]. Cry5B has also been shown to have synergistic whole-worm effects when combined with the cholinergic anthelmintics levamisole. Combination of Cry5B and levamisole results in enhanced clearing of parasites when compared to any of the single compounds when used alone [18, 19]. The synergistic interaction between the cholinergic anthelmintics and the effects of Cry5B has been described at the whole-worm level, but the location, mechanism and time-course of the interaction is not known.

*Ascaris suum* appears genetically and phenotypically to be the same species as *Ascaris lumbricoides* of humans [20]. Here we utilize a Ca^2+^ imaging protocol adapted for *Ascaris suum* enterocytes and histology on the intestine of this nematode to study the effects of Cry5B and levamisole. We observed that Cry5B had concentration-dependent and time-dependent effects on the amplitude of the Ca^2+^ signal that can be directly related to damage of the intestine. Destruction of the intestine tissue by Cry5B occurred within 6 hours. We explored the effects of combining Cry5B and levamisole on the Ca^2+^ signal and tissue integrity: the result of the combination was a faster and larger Cry5B induced Ca^2+^ signal than when it was applied by itself. We identified the intestine as a shared site of action for the cholinomimetic anthelmintic levamisole and activated Cry5B; and we observed that the time-course of the interaction was under 2 hours.

## Materials and Methods

### Collection and maintenance of A. suum worms

Adult female *A. suum* worms were collected from the JBS Swift and Co. pork processing plant at Marshalltown, Iowa. Worms were maintained in *Ascaris* Ringers Solution (ARS: 13 mM NaCl, 9 mM CaCl_2_, 7 mM MgCl_2_, 12 mM C_4_H_11_NO_3_/Tris, 99 mM NaC_2_H_3_O_2_, 19 mM KCl and 5 mM glucose pH 7.8) at 32 °C for 24 hours to allow for acclimatization before use in experiments. The solution was changed daily, and worms were used within three days for experiments. All the worms were examined at the start of each day and removed if they were damaged or immotile.

### A. suum cDNA synthesis and RT-PCR detection of Cry5B mRNA

Dissection of body wall and intestinal tissue was conducted on adult *A. suum* females as previously described [21, 22]. The body wall and intestinal tissue were homogenized separately in 1 ml of Trizol reagent using a mortar and pestle, followed by total RNA extraction according to the Trizol Reagent protocol (Life Technologies, USA). One microgram (1 µg) of total RNA from each tissue was used to generate cDNA by reverse transcription (RT) using SuperScript IV VILO^TM^ Master Mix (Life Technologies, USA) following the manufacturer’s protocol. PCR was conducted to detect the presence of *Asu-bre-5* and *Asu-cdh-8* using primers targeting coding regions of each gene (Supplementary Table S1)*. Asu-gapdh* was used as a reference gene. The negative controls used no cDNA template. The PCR conditions were an initial denaturation for 2 min at 98 °C, followed by 35 cycles of 98 °C for 30 sec, 58 °C for 35 sec, 72 °C for 45 sec, and a final extension at 72 °C for 10 min using GoTaq^®^ G2 Hot Start Green Master Mix (Promega, USA). The PCR products of each gene were then separated on 2% agarose gels containing SYBR^®^ Safe DNA Gel Stain (ThermoFisher Scientific), at 100V, followed by visualization under UV light to confirm the presence of the genes. All photographs were acquired using Visionworks^TM^ software (Analytik Jena) with an exposure setting of 3 seconds per 1 frame. Original gel pictures are presented in Supplementary Fig. S1.

### Analysis of mRNA levels by Quantitative Real-time PCR

We quantified mRNA transcript levels of *Asu-bre-5* and *Asu-cdh-8* in the intestine and body wall of adult female *A. suum*. Amplicons ranging from 150 to 200 bp were generated by qPCR from each cDNA sample in triplicate. *Asu-gapdh* was used as the reference gene (for qPCR primers see Supplementary Table S2). The quantitative PCR reaction mixture consisted of 1 μl of cDNA template, 1 μl of the forward and reverse primer, and 10 μl of PowerUp^TM^ SYBR^TM^ Green Master Mix (Applied Biosystems, ThermoFisher, USA), with the final volume made up to 20 μl with nuclease-free water. The thermocycler conditions included an initial denaturation for 10 seconds at 98°C, 40 cycles of 98°C for 15 seconds and 58°C for 30 seconds followed by a final melting curve step. Cycling was performed using a QuantStudio^TM^ 3-96 well 0.1 mL Block Real time PCR Detection system (ThermoFisher, USA), and transcript quantities were derived by the system software, using the generated standard curves. mRNA expression levels for each gene (*Asu-bre-5 and Asu-cdh-8*) were estimated relative to the reference gene (*Asu-gapdh*) using the Pfaffl Method [23]. The qPCR experiments were repeated 3 times for each gene (all subunit mRNA quantifications were performed in triplicate for each worm’s body wall sample and intestinal tissue sample: 15 biological replicates each with three technical replicates).

### Preparation and loading Fluo-3AM

Intestinal tissues were loaded with Fluo-3AM as previously described [22]. Briefly, a 2 cm section of the intestine was removed from the body piece using fine forceps and cut open. The intestinal flap was placed in a Warner RC26G recording chamber (Warner Instruments, Hamden, CT)) and immobilized using a 26 x 1mm x 1.5mm grid slice anchor (Warner Instruments, Hamden, CT), bathed in *Ascaris* Perienteric Fluid APF (23 mM NaCl, 110 mM NaAc, 24 mM KCl, 1 mM CaCl_2_, 5 mM MgCl_2_, 5 mM HEPES, 11 mM D-glucose)(Fig. 1A). The chamber temperature was maintained at 34-36 °C using a Dual Automatic Temperature Controller and inline heater (Warner Instruments, Hamden, CT) maintained at Fluo-3AM loading was achieved by incubating the intestine in APF solution with no added CaCl_2_ ([Ca^2+]^]<100 µM) containing 5 µM Fluo-3AM and 10% Pluronic F-127 (10% v/v) for one hour.. After incubation, the Fluo-3AM solution was discarded, and the sample was incubated in APF containing 1 mM CaCl_2_ for an additional 15 minutes at 34-36°C to promote Ca^2+^ loading (Fig. 1B). All incubations were done in the absence of light to prevent degradation of the fluorescent dye. At the start of all experiments, intestinal preparations were left under blue light for a minimum of 3 minutes to promote settling and equilibration of the fluorescent signal and monitor for any spontaneous Ca^2+^ signals before application of any compound. For each recording the APF was removed using an exhaust line and the sample was exposed to either fresh 1 mM CaCl_2_ APF, 100 µg/ml Cry5B, 10 µg/ml Cry5B or 100 µg/ml Cry5B and 100 mM galactose for 6 hours or 10 µg/ml Cry5B, 30 µM levamisole, 10 µg/ml Cry5B and 30 µM levamisole for 2 hours. The solution level of the chamber was constantly maintained throughout the recordings. The temperature of the chamber was maintained between 34-36°C for the entire recording, any samples where the temperature deviated from this range were discarded. All solutions were delivered to the chamber using transfer pipettes away from the intestinal preparation.

**Figure 1:**
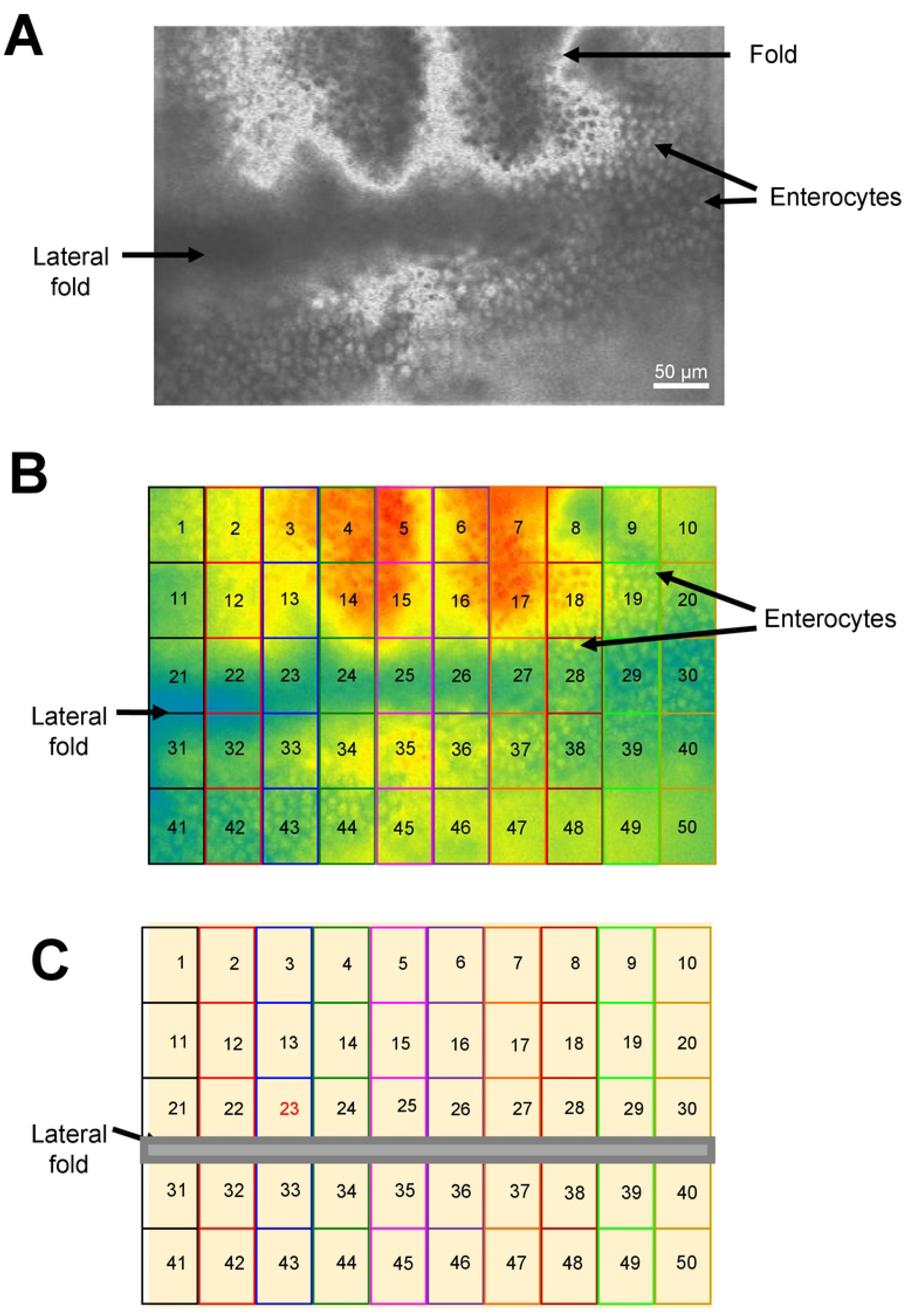
Illustration of intestinal regions: A) Photograph of intestinal preparation under white light. Key structures are highlighted. B) Photograph of the same intestinal preparation shown in (A) under blue light after Fluo-3AM treatment and Ca^2+^ loading, highlighted with the 50 regions used to record Ca^2+^ signals. C) Cartoon illustration of (B) highlighting the 50 regions used to record from a signal intestinal piece. Lateral line is included.

### Measurement of Ca^2^ fluorescence

All recordings were performed with a Nikon Eclipse TE300 microscope (20X/0.45 Nikon PlanFluor objective), fitted with a Photometrics Retiga R1 camera (Photometrics, Surrey, BC, Canada). Light control was achieved using a Lambda 10-2 two filter wheel system with a shutter controller (Lambda Instruments, Switzerland). Filter wheel one was set on a green filter (510-560 nM bandpass, Nikon USA) between the microscope and camera. Filter wheel two set on the blue filter (460-500 nM, bandpass, Nikon, USA) between a Lambda LS Xenon bulb light box which delivered light via a fiber optic cable to the microscope (Lambda Instruments, Switzerland). The blue light emission was controlled using a shutter. All Ca^2+^ signal recordings were acquired and analyzed using MetaFluor 7.10.2 (MDS Analytical Technologies, Sunnyvale, CA). Exposure times were 150ms with 2x binning. Maximal Ca^2+^ signal amplitudes (ΔF/F_0_ %) for all stimuli applied were calculated using the equation F-F0/F0 where F is the fluorescent value and F0 is the baseline fluorescent value, which was determined as the lowest value before the largest rise in fluorescence for all recordings analyzed. Representative traces were generated using the same formula, with F0 being the value before a detectable increase in the fluorescence. For the 1 mM CaCl_2_ bath solution control experiments (Fig. 3A & C) F0 was determined to be the value before the largest rise in Ca^2+^. For the control CaCl_2_ responses (Fig 3) F0 was determined as the value before stimulus application. Rise times were calculated by normalizing the trace during stimulus exposure, with the lowest fluorescence value being represented by 0% and the highest being 100%. The peak time was calculated by subtracting the time when the stimulus was applied from the time the signal reached 100%.

### Regions used for measurements of the Ca^2+^ signals and % response area

Ca^2+^ signals from each intestine were collected from 50 squares each 50 µm x 50 µm from rectangular 125,000 µm^2^ areas of the intestine that included 800-1000 enterocytes (Fig. 1B & C). The relative fluorescence amplitude was determined and followed over time for each of the individual 50 square regions. Long-term control preparations were not exposed to any test agents and were incubated in 1 mm CaCl_2_ APF and followed over 6 hours. A 10 mM CaCl_2_ test pulse was added to the chamber as a test of viability of the preparation. An increase >10% in relative Fluo-3 fluorescence to the 10 mM CaCl_2_ pulse was taken as an indication of the positive health of the preparation: preparations were rejected if the responses were <10% as not being viable. For the 6-hour incubations, intestines were exposed to either 10 µg/ml Cry5B, 100 µg/ml Cry5B, or 100 µg/ml Cry5B and 100 mM galactose. For the 2-hour recordings preparations were exposed to either 10 µg/ml Cry5B, 30 µM levamisole as the sole active agent or a combination of 10 µg/ml Cry5B and 30 µM levamisole. Any of the 50 µm x 50 µm regions whose Ca^2+^ amplitude responses to the anthelmintic stimulus that was smaller than the average amplitude of the spontaneous Ca^2+^ signals (2.4% ±0.1% Fig. 2C) were discarded. The reason for their rejection was that we could not rule out the possibility that these signals were, themselves, spontaneous rather than produced by the anthelmintic stimulus.

**Figure 2:**
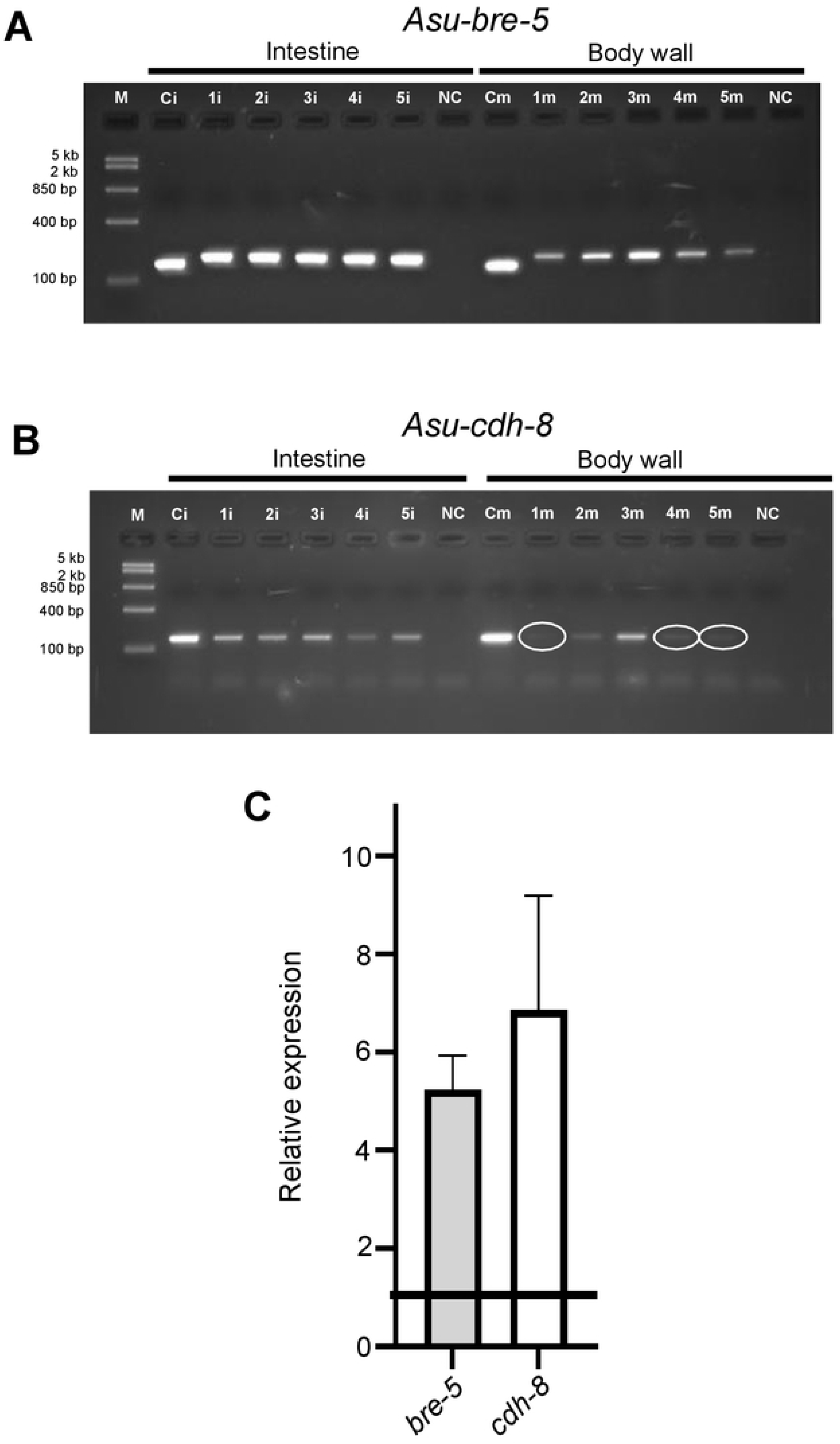
Cry5B targets are present in *Ascaris suum* intestine: RT-PCR analysis of intestine (1i, 2i, 3i, 4i, 5i) and body wall (1m, 2m, 3m, 4m, 5m) of five separate female *A. suum* worms. Each lane represents the intestine or body wall of an individual worm. *Asu-gapdh* from the intestine (Ci) and body wall (Cm) was used as a positive control. N.C. = no template negative control. M = Fast Ruler Middle Range DNA Ladder (TermoFisher Scientific). A) *Asu-bre-5* B) *Asu-cdh-8.* Note reduced intensity of bands highlighted by white circles in the body wall samples. All images are cropped. Images were taken under UV light with an exposure setting of 3 s per 1 frame. Original gel images are presented in Supplementary Fig. S1. C) Differential expression of Cry5B targets between intestine and body wall. Bar chart (expressed as mean ±SEM) demonstrating transcript level analysis from intestinal samples for *bre-5* (grey bar) (5.24 ± 0.68) and *cdh-8* (white bar) (6.86 ± 2.32) when compared to paired body wall samples. n = 15 individual worms over three biological replicates.

### Histology

Sections of *Ascaris* body were cut along the lateral line and opened, exposing the intestine. For 6-hour comparisons, sections were incubated in either: APF for 1, 2, or 6 hours; APF containing 100 µg/ml Cry5B for 1, 2, or 6 hours; APF containing 10 µg/ml Cry5B for 1, 2, or 6 hours; or APF containing 100 µg/ml Cry5B and 100 mM galactose for 1, 2, or 6 hours. For the two-hour incubations samples were incubated in: APF containing 30 µM levamisole for 2 hours; or APF containing 10 µg/ml Cry5B and 30 µM levamisole for 2 hours.

Following treatment, the segments of adult *Ascaris* worms were preserved in 10% buffered neutral formalin. Preserved specimens were placed in histology cassettes for paraffin embedding and sectioning. Hematoxylin and eosin staining was performed on 5 µm slides [24]. Sections were viewed and images were captured on an Olympus BX60 microscope fitted with an Olympus DP70 camera (Olympus, Tokyo, Japan). Image captures were achieved using Olympus DP controller software (Olympus, Tokyo, Japan) under 20x magnification and exposure settings of 1/350 seconds per frame.

### Preparation of Cry5B

Cry5B was supplied by Raffi Aroian in both active and inactive form. Inactive Cry5B was activated using elastase at a Cry5B to elastase mass ratio of 200:1, overnight at room temperature [25]. Active Cry5B was stored at −80°C suspended in 20 mM HEPES (pH 8.0). Cry5B solutions were made fresh for each experiment.

### Statistical Analysis

Statistical analysis of all data was made using GraphPad Prism 9.0 (Graphpad Software, Inc., La Jolla, CA, USA). We repeated our experiments to ensure reproducibility. The total number of female worm intestinal preparations, the total number of regions showing responses, the concentrations, and durations of applications are provided in the figure legends of the figures. Analysis of Ca^2+^ amplitudes and time to peak were made using either unpaired for separate preparations or paired when the responses in the same preparation were being followed using student *t*-tests *P <* 0.05. The *t*-test (paired or unpaired) used is stated in the figure legends.

### Chemicals

Source of chemicals: Cry5B was provided by Raffi Aroian. Galactose, levamisole and pyrantel were procured from Sigma Aldrich. All other chemicals were supplied by Fisher Scientific.

### Data availability statement

The datasets analyzed during the current study are available in Wormbase Parasite, The European Nucleotide Archive (ENA) and UNIPROT repositories, https://parasite.wormbase.org/index.html, www.elixir-europe.org/platforms/data/core-data-resources, www.uniprot.org. Links to the datasets are presented in supplementary Table 3.

## Results

### Cry5B targets are present in Ascaris suum intestine

The *Bacillus thuringiensis* (Bt) bacteria produce different pore forming toxins referred to as Crystal 3D or Cry proteins. Cry5B is a Cry protein that is nematocidal and is understood to have a similar mode of action as the insecticidal Cry proteins. Previous studies have described receptors for Cry5B that are: 1) glycosphingolipids that are produced by the acetylglucosaminyltransferase activity of BRE-5 [15, 16] and; 2) the cadherin CDH-8 which are responsible for oligomerization of Cry5B [17]. Loss of both genes results in a level of resistance to Cry5B in *C. elegans*.

To determine if BRE-5 and CDH-8 are present in *Ascaris suum,* we blasted the *C. elegans* orthologues against the *Ascaris suum* genome and identified the conserved genes, *Asu-bre-5* and *Asu-cdh-8* (Supplementary Table 3). To determine if message for these genes is expressed in the intestine and/or body wall, we generated cDNA from RNA pools from the intestine and body wall and screened using primers targeting *Asu-bre-5* and *Asu-cdh-8*, Fig. 2A & B (Supplementary Table 1). Fig. 2 shows that *Asu-bre-5,* Fig. 2A, and *Asu-cdh-8,* Fig. 2B, were present in both the intestine and body wall in all pools tested. The intensity of the bands in the intestine was higher than the body wall for both *bre-5* and *cdh-8*.

To determine if expression of *bre-5* and *cdh-8* were higher in the intestinal tissue, we performed qPCR comparing the relative expression level between the body wall and the intestine. Fig. 2C shows that the relative expression levels of *Asu-bre-5* and *Asu-cdh-8* were higher in the intestine than the body wall (5.2-fold for *bre-5* and 6.9-fold for *cdh-8*). These observations suggest that the Cry5B targets are conserved in *Ascaris suum* and that these targets have a higher level of expression in the intestine, but the targets are also present in the body wall of the parasite.

### Ca^2+^ signaling and effects of Cry5B on the intestine

With message indicating the presence of glycosphingolipids receptors produced by BRE-5 activity and cadherin CDH-8 receptors, we sought to determine the time course and nature of the effects of Cry5B exposure on the *Ascaris* intestine. We hypothesized that Cry5B would increase Ca^2+^ entry into the enterocytes, a feature seen with other pore forming toxins including phobalysin from *Photobacterium dameselae* [26]. Although, the effects of Cry5B have been studied on whole animals, the timing of direct effects on the enterocytes of the parasites were not known.

In control experiments, we found that we could maintain viable intestine tissues and perform Ca^2+^ imaging for at least 6 hours, Fig. 3A. We observed a slow initial exponential decline in the relative Ca^2+^ fluorescence during the first two hours, a stable signal thereafter except for the occasional small spontaneous transient signals. We analyzed the largest transient signals during the 6-hour period and found the average amplitude of the spontaneous Ca^2+^ fluctuations were 2.4% ± 0.1%, Fig. 3B; white bar. Although the spontaneous fluctuations indicate a healthy and viable tissue, as a positive control, we transiently increased the Ca^2+^ concentration by adding 10 mM CaCl_2_ to the bath after the 6-hour recording. As seen in Figs. 3A & C, the pulse of 10 mM CaCl_2_ produced in a large increase in relative fluorescence that had a mean of 40.36% ± 0.88%, Fig. 3B, red bar. When the Ca^2+^ was returned to 1 mM, the signal returned to the baseline, indicating that the tissue was not compromised. In damaged or deteriorating intestine preparations, the 10 mM CaCl_2_ test did not produce a Ca^2+^ signal increase nor a change on washing with 1mM CaCl_2_ in APF.

**Figure 3:**
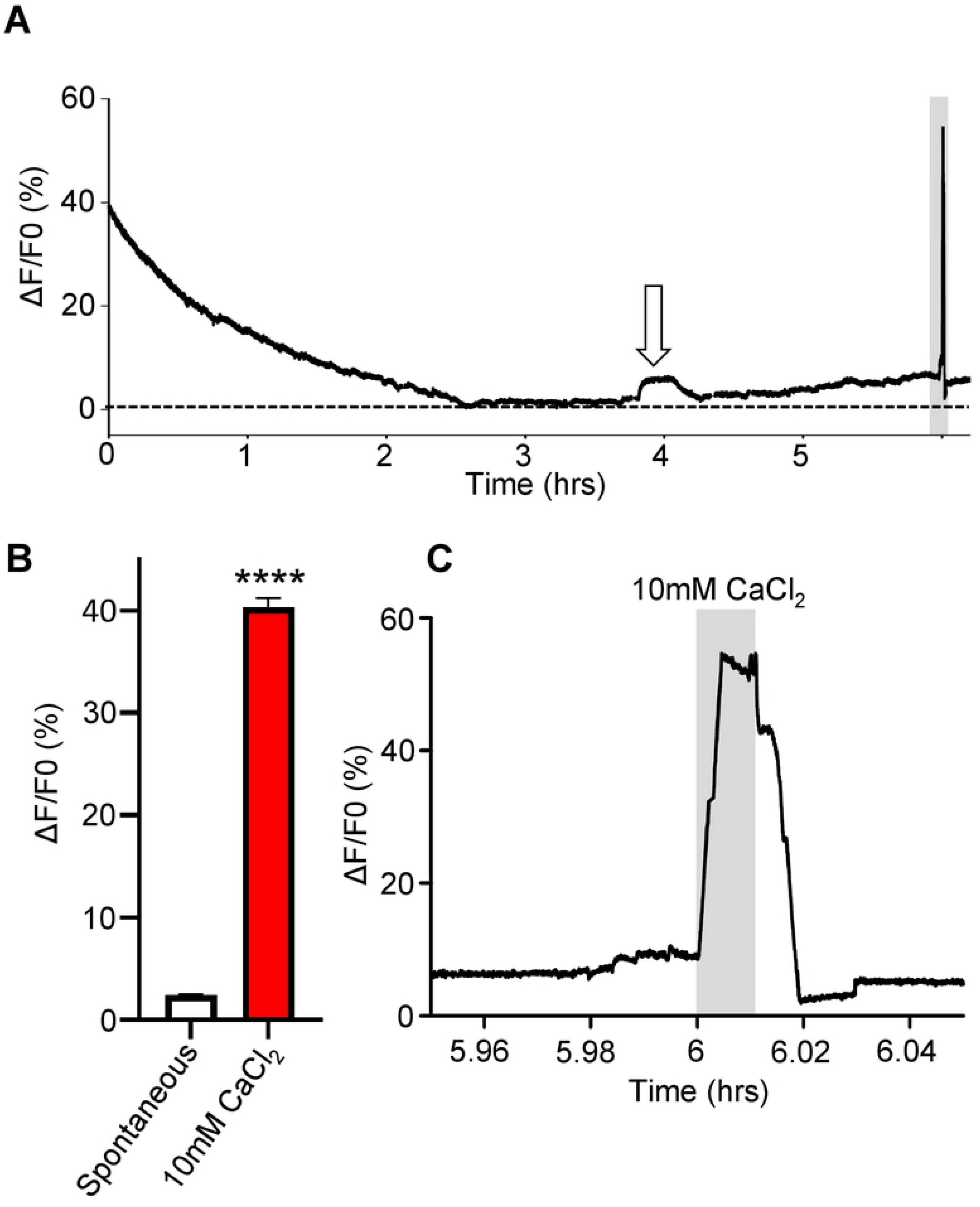
Long-term Ca^2+^ imaging: A) Representative trace of a 6-hour Ca^2+^ imaging recording on *Ascaris suum* intestines in the presence of APF. White arrow highlights a spontaneous Ca^2+^ signal. Grey box indicates application of 10 mM CaCl_2_ a positive control. B) Amplitudes of spontaneous Ca^2+^ signals (white bar) and control 10 mM CaCl_2_ amplitudes (red bar). **** Significantly different to spontaneous. Spontaneous vs 10 mM CaCl_2_ *P* <0.0001 *t* = 41.76, *df* = 249, paired *t*-test. C) Close up of 10 mM CaCl_2_ signal from the end of the trace in A. Grey box highlights 10 mM CaCl_2_ application. *n* = 5 individual female intestines with a total of 250/250 regions providing viable responses which were used to generate the mean and SEM for B.

Initially, we tested a concentration of 100 µg/ml Cry5B on the intestine, based on whole animal *in vitro* studies on *Ascaris suum* [13, 27]. These studies reported that 100 µg/ml Cry5B was the *EC_50_*concentration for *Ascaris suum* being killed before 24 hours. Application of Cry5B to the intestines produced no immediate effects on the Ca^2+^ signal, Fig. 4A. After one hour we observed small fluctuations in the signals which were followed by larger increases in fluorescence with an average maximal amplitude of 45.73% ± 2.8%, Fig. 4D: red bar. The peak Ca^2+^ increases, which lasted around two hours, were seen at an average of 80 mins ± 3.3 mins, Fig. 4E: red bar. After the peak, the Ca^2+^ signal declined slowly in a jerky manner below the zero level as the integrity of the enterocytes and intensity of the Fluo-3 fluorescence was lost which was observable down the microscope. The large Cry5B Ca^2+^ peak responses occurred in most of the 50 regions of the recorded intestines, 94.8% ± 3.2%, Fig. 4F: red bar, and suggests that the distribution of the Cry5B receptors is widely and evenly distributed over the area of the intestine. Thus we observed that the Ca^2+^ signals on the intestine of 100 µg/ml Cry5B are slow in onset (average peak 80 mins) but they are bigger than the levamisole and diethylcarbamazine anthelmintic compounds that we have tested previously [21, 22].

**Figure 4:**
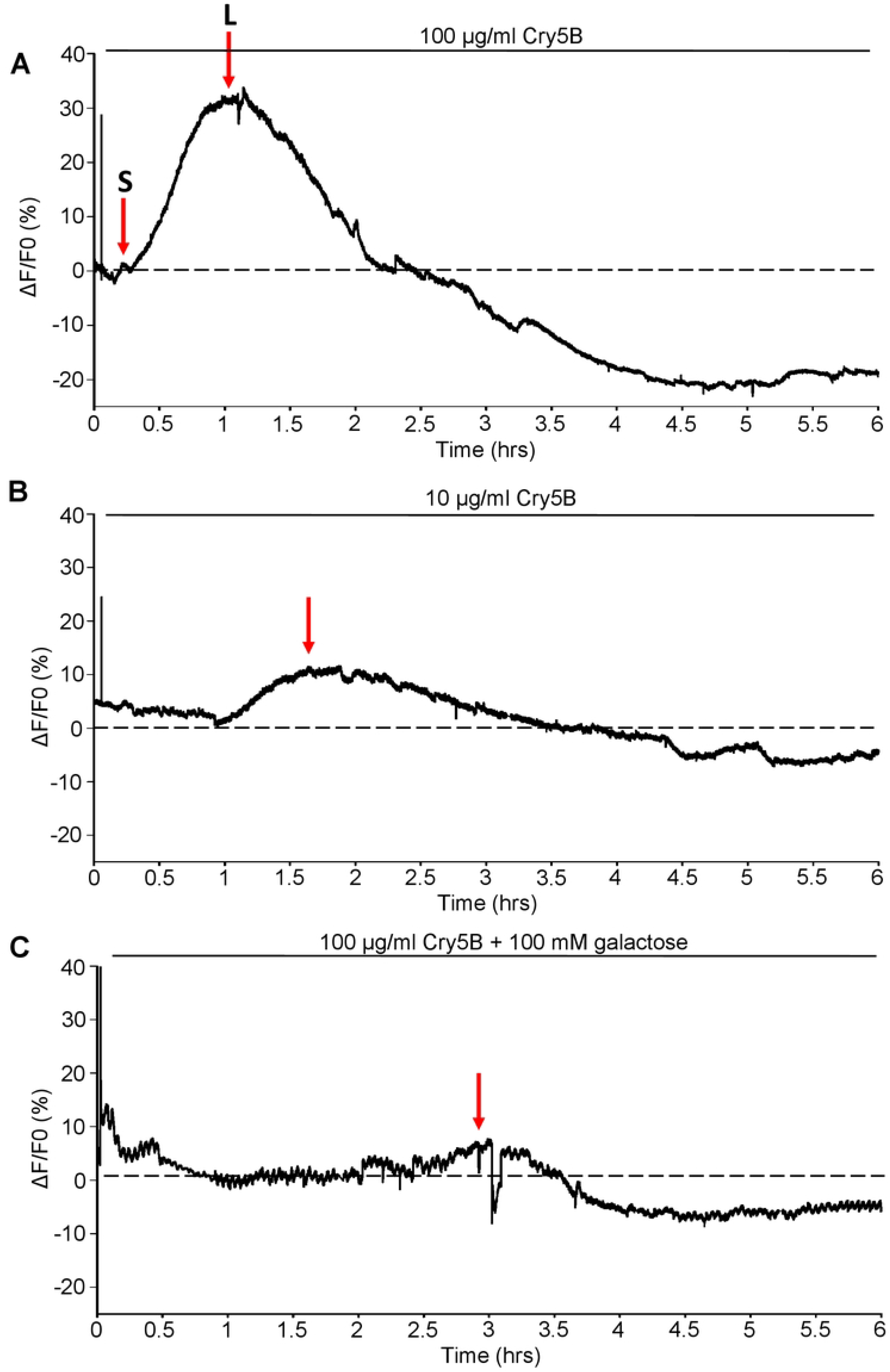
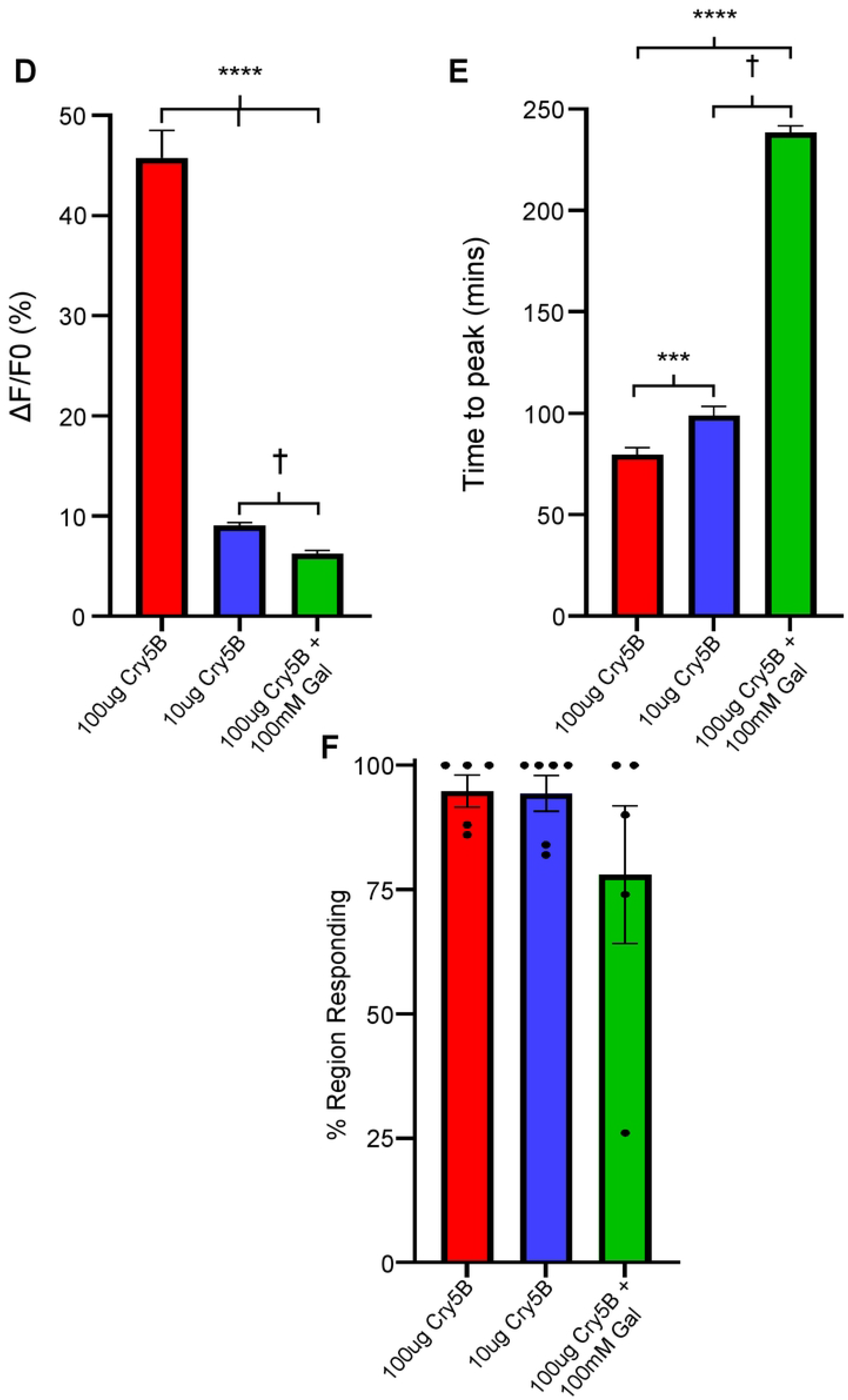
Cry5B stimulates concentration dependent Ca^2+^ responses on *A. suum* intestines that are inhibited by galactose. A) Representative trace of a 6-hour Ca^2+^ recording to 100 µg/ml Cry5B. Dotted line highlights 0% F/F_0_. **S** indicates initial small increase in Ca^2+^ and **L** represents the secondary larger response B) Representative trace of a 6-hour Ca^2+^ recording to 10 µg/ml Cry5B. Dotted line highlights 0% F/F_0_. The red arrow indicates the peak of the Ca^2+^ signal. C) Representative trace of a 6-hour Ca^2+^ recording to 100 µg/ml Cry5B + 100 mM galactose. Dotted line highlights 0% F/F_0_. The red arrow indicates the peak of the Ca^2+^ signal. D) Average maximal amplitudes of Ca^2+^ fluorescence over 6 hours in response to 100 µg/ml Cry5B treated (red bar), 10 µg/ml Cry5B treated (blue bar) and 100 µg/ml Cry5B + 100 mM galactose treated (green bar). **** Significantly different to 100 µg/ml Cry5B. (100 µg/ml Cry5B vs 10 µg/ml Cry5B, *P* = <0.0001 *t* = 14.39, *df* = 518, unpaired *t*-test, 100 µg/ml Cry5B vs 100 µg/ml Cry5B + 100 mM galactose *P* = <0.0001 *t* = 14.55, *df* = 485, unpaired *t*-test). † Significantly different to 10 µg/ml Cry5B (10 µg/ml Cry5B vs 100 µg/ml Cry5B + 100 mM galactose *P* = <0.0001 *t* = 6.437, *df* = 531, unpaired *t*-test. E) Average time to maximal peak amplitude over 6 hours in response to 100 µg/ml Cry5B treated (red bar), 10 µg/ml Cry5B treated (blue bar) and 100 µg/ml Cry5B + 100 mM galactose treated (green bar). *** Significantly different to 100 µg/ml Cry5B. (100 µg/ml Cry5B vs 10 µg/ml Cry5B, *P* = <0.0007 *t* = 3.412, *df* = 518, unpaired *t*-test. **** Significantly different to 100 µg/ml Cry5B. 100 µg/ml Cry5B vs 100 µg/ml Cry5B + 100 mM galactose *P* = <0.0001 *t* = 14.55, *df* = 485, unpaired *t*-test). † Significantly different to 10 µg/ml Cry5B (10 µg/ml Cry5B vs 100 µg/ml Cry5B + 100 mM galactose *P* = <0.0001 *t* = 6.437, *df* = 531, unpaired *t*-test). F) Average percent of regions showing a Ca^2+^ response larger than the average spontaneous amplitude in untreated intestines in 100 µg/ml Cry5B treated (red bar), 10 µg/ml Cry5B treated (blue bar) and 100 µg/ml Cry5B + 100 mM galactose treated (green bar). N.S. not significant compared to 100 µg/ml Cry5B (100 µg/ml Cry5B vs 10 µg/ml Cry5B, *P* = <0.9264 *t* = 0.095, *df* = 9, unpaired *t*-test, 100 µg/ml Cry5B vs 100 µg/ml Cry5B + 100 mM galactose *P* = <0.271 *t* = 1.183, *df* = 8, unpaired *t*-test). 100 µg/ml Cry5B *n* = 5 individual female intestines with a total of 237/250 regions providing viable responses which were used to generate the mean and SEM for D & E. Note: 13/250 regions showed no response. 10 µg/ml Cry5B *n* = 6 individual female intestines with a total of 283/300 regions providing viable responses which were used to generate the mean and SEM for D & E. Note: 17/300 regions showed no response. 100 µg/ml Cry5B + 100 mM galactose *n* = 5 individual female intestines with a total of 195/250 regions providing viable responses which were used to generate the mean and SEM for D & E. Note: 55/250 regions showed no response.

### Histology

With Cry5B producing large Ca^2+^ signals in the intestine, we wanted to see how the Ca^2+^ signal correlated with histological effects on the intestine. We incubated body *Ascaris* flap preparations that were cut along one lateral line through into the intestine in APF (untreated) or in APF containing 100 µg/ml Cry5B. We used the body flaps with the attached intestine to maintain the intestine position and to prevent folding. At the 2-hour time, where we observed the biggest Ca^2+^ signal amplitude, the sections in the Cry5B treated sample showed widespread evidence of vacuolation throughout the length of the enterocyte, and signs of necrosis of the enterocytes, Fig. 5A: right panel. The untreated control sections at 2-hours, Fig. 5A: left panel, did show some round vacuoles at the apical surface of the enterocytes, but the individual enterocytes did not show initial signs of separation between themselves and the vacuolation was more limited. The differences between the treated and untreated 2-hour sections were noticeable. At 6-hours, we observed nearly complete disruption of the enterocytes and separation from the basolateral membrane when treated with 100 mg/ml Cry5B (Fig. 5B: right panel) with no detection of the brush border or intact enterocytes. The untreated 6-hour section showed increased vacuolation, but the brush boarder was still intact, and enterocytes were not detached from the basolateral membrane, Fig. 5B: left panel. Taken together, the histology demonstrates a direct effect of activated Cry5B on the intestine and the damage on this tissue that correlates with the Ca^2+^ signal. We have been able to establish a time frame of the 100mg/ml Cry5B action that starts at around 1 hour and results in destruction of the integrity of the enterocytes in less than 6 hours. There is loss of the brush boarder, significant vacuolation and separation of adjacent enterocytes from the basolateral border and necrosis and cell death of the enterocytes that will damage the barrier properties of the intestine.

**Figure 5:**
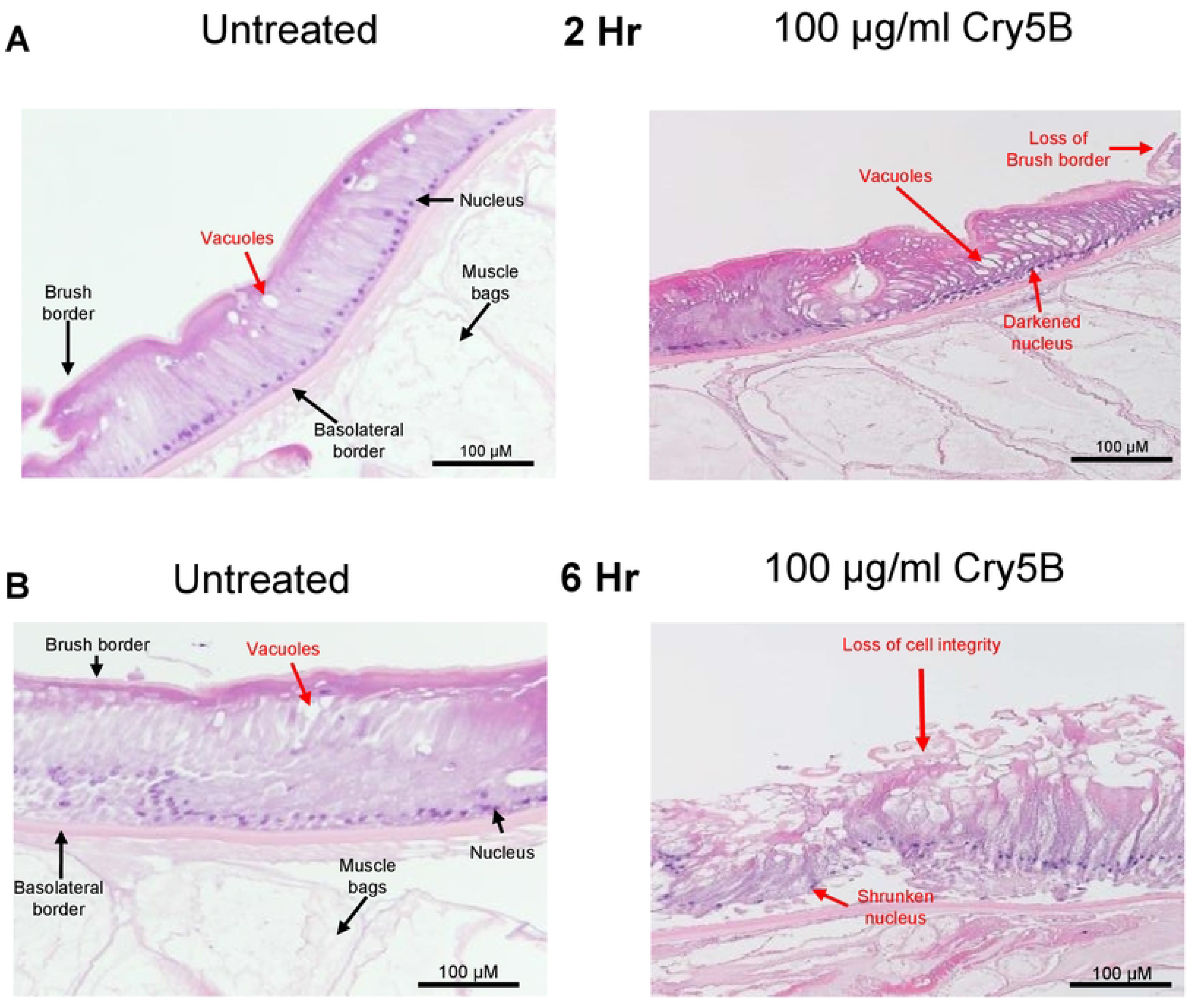
100 µg/ml Cry5B causes severe damage to the intestine of *A. suum* over 6 hours. Transverse H & E sections of the *Ascaris* intestine and underlying muscle bags in untreated (APF) *left column* and 100 µg/ml Cry5B treated *right column* at different time courses. A) 2hrs of treatment and B) after 6hrs of treatment. Major structures and evidence of tissue damage are highlighted. Note that the nuclei of the Cry5B treated preparations are shrunken and darker.

### Concentration-dependent effects of Cry5B

We have seen that 100 µg/ml Cry5B causes a large slow transient increase in the Ca^2+^ signal and histological damage to the intestine. Urban et al. 2013 showed that low concentrations of Cry5B can still kill parasites but the incubation times are longer. We tested a concentration of 10 µg/ml Cry5B, Fig. 4B & D; blue bar. The lower concentrations of Cry5B, still initiated Ca^2+^ signals, but they were significantly smaller than responses produced by 100 µg/ml Cry5B. The average amplitude of the 10 µg/ml Cry5B signals was 9.05% ± 0.3%. The time to peak for 10 µg/ml Cry5B treated samples was 99 mins ± 4.4 mins, Fig. 4E: blue bar, on average and was significantly slower than that of 100 µg/ml Cry5B. Again, after the peak the Ca^2+^ signal decreased below the zero level. Although the times to peaks were on average smaller and were slower to develop for 10 µg/ml Cry5B, we observed that the effects were still widespread and covered 94.3% ± 3.6% of the 50 observed regions in the different preparations during the recording, Fig. 4F: blue bar.

We tested the effects of 10 µg/ml Cry5B for the 2-hour and the 6-hour on the histology sections of the intestines and observed changes that were less severe. At the 2-hour time point nearer the peak of the Ca^2+^ signal, we observed differences between the untreated and 10 µg/ml Cry5B treated preparations, Fig 6A; right panel. There were larger elongated vacuoles occurring in the Cry5B treated samples near the apical region of the enterocytes and some separation between adjacent enterocytes: these sections contrasted with those of the 2-hour untreated sections, Fig. 6A; left panel, which showed no separation between enterocytes and smaller round vacuoles. The 6-hour 10 µg/ml Cry5B sections had more vacuoles which were larger in the treated sections compared to the untreated sections; and there was some erosion of parts of the brush border, Fig 6B. We concluded that the effects of Cry5B on Ca^2+^ signaling and histology on the intestine were concentration-dependent, with 10 µg/ml Cry5B producing smaller and slower Ca^2+^ peak amplitudes with damage occurring at a slower rate than with the 100 µg/ml. However, both 10 µg/ml and 100 µg/ml had effects on nearly all the 50 regions of the intestines; and the 6-hour time is sufficient for activated toxin to damage to the entire intestinal tissue.

**Figure 6:**
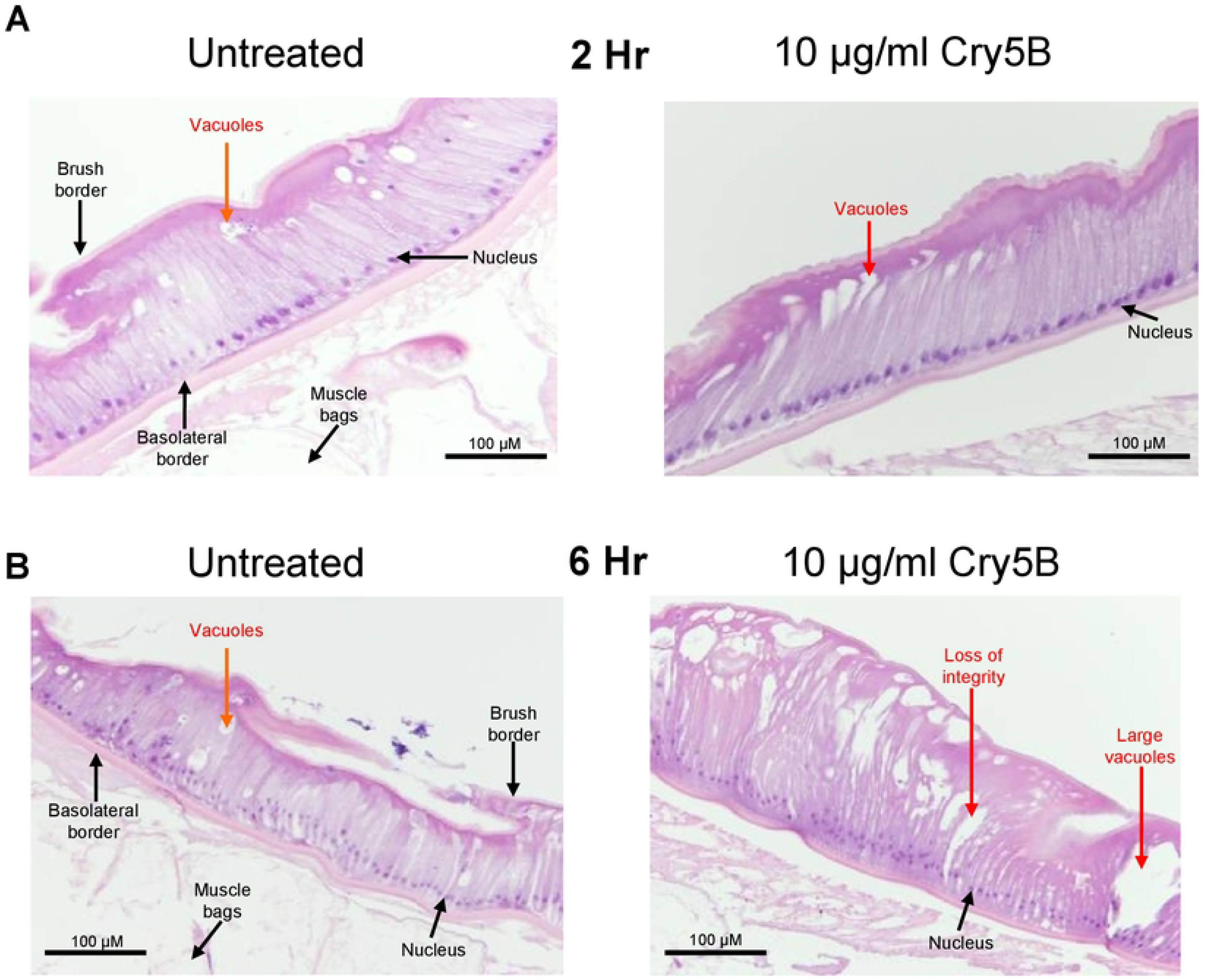
10 µg/ml Cry5B causes damage to the intestine of *A. suum* over 6 hours. Transverse H & E sections of the *Ascaris* intestine and overlying muscle bags in untreated (APF) *left column* and 10 µg/ml Cry5B treated *right column* at: A) after 2hrs B) 6hrs. of treatment. Major structures and evidence of tissue damage are highlighted. Note the increased number of vacuoles in the Cry5B treated sample at 6 hrs. and loss of cellular integrity compared to untreated.

### Galactose inhibits the action of Cry5B

The action of Cry5B on nematodes has included interaction with glycosphingolipid receptors produced by BRE-5 [9, 15, 16, 25, 28] that 100 mM galactose inhibits the action of Cry5B by competing for binding to the glycosphingolipid and promotes survivability long-term. We have identified the *Ascaris suum* orthologue for *bre-5* is present in the intestine, Fig. 2A, suggesting that the observed Cry5B stimulated Ca^2+^ signal and subsequent intestinal damage involves interaction with BRE-5 produced glycosphingolipids. We tested the effects of 100 mM galactose on the effects of 100 µg/ml Cry5B on the Ca^2+^ signal and histology of the intestine.

100 µg/ml galactose strikingly inhibited in the Ca^2+^ signal produced by Cry5B, Figs. 4A & C. The increase in the 100 µg/ml Cry5B Ca^2+^ signal in the presence of 100 mM galactose was significantly smaller than either the 100 µg/ml Cry5B or the 10 µg/ml Cry5B, Fig. 4D: green bar. Additionally, the average time to peak was significantly longer than either the 100 µg/ml Cry5B and 10 µg/ml Cry5B with average rise time being 238 mins ± 3.4 mins, Fig. 4E: green bar. Although the average percentage of the 50 regions in the different preparations showing a Ca^2+^ responses larger than the spontaneous signals of the untreated intestines was reduced to 78% ± 13.84, Fig. 4F: green bar, it did not reach statistical significance. Histology showed that in the presence of 100 mM galactose the Cry5B mediated damage to the intestines was significantly inhibited, even at 6-hours, Fig. 7A. These observations suggest that the mode of action of Cry5B is conserved in the *Ascaris suum* intestine and that Cry5B initiates a strong Ca^2+^ signal and intestinal damage that starts from the interaction with glycosphingolipid receptors. The competitive binding by galactose limits intestinal damage.

**Figure 7:**
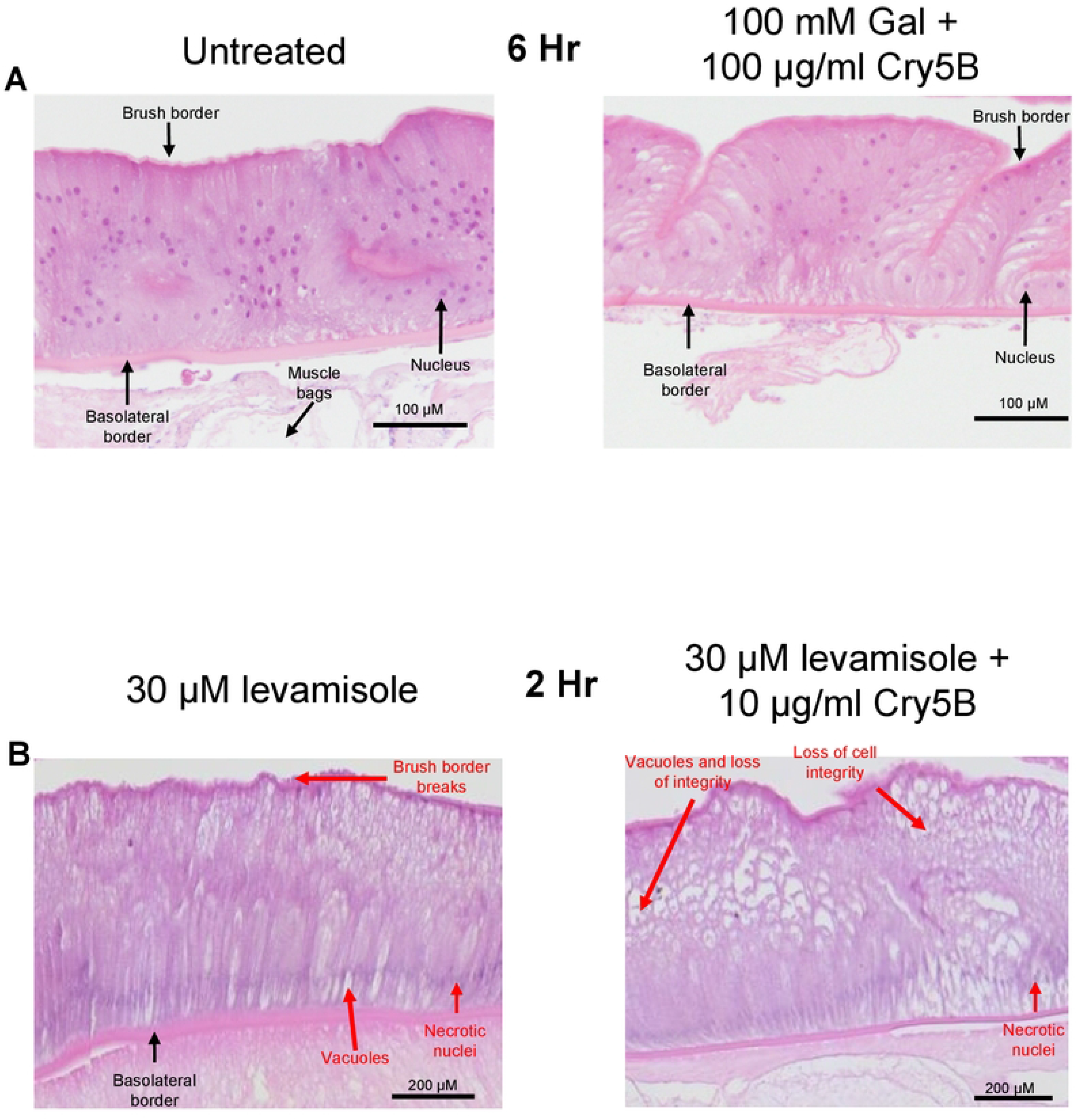
Histological effects of 100 mM galactose and levamisole on Cry5B. A) 100 mM galactose prevents intestinal damage caused by 100 µg/ml Cry5B. Transverse H & E sections of the *Ascaris* intestine and underlying muscle bags in untreated (APF) *left column* and 100 µg/ml Cry5B + 100 mM galactose treated *right column* after 6hrs. of treatment. Major structures are highlighted. No indications of major damage including loss of cell integrity or loss of brush border was observed in either preparation, apart from a few vacuoles. Note: loss of attached muscle cell bags in 100 µg/ml Cry5B + 100 mM galactose treated example is due to mechanical damage during preparation. B) Transverse H & E sections of the *Ascaris* intestine and underlying muscle bags after 2 hours in 30 µM levamisole (left panel) and 30 µM levamisole + 10 µg/ml Cry5B (right panel). Major structures and evidence of tissue damage are highlighted. Treatment with both levamisole alone and levamisole + Cry5B resulted in the loss of recognizable nuclei. Addition of 10 µg/ml Cry5B and 30 µM levamisole resulted in significant loss of cell integrity compared to 30 µM levamisole or 10 µg/ml Cry5B alone.

### Levamisole potentiates Cry5B Ca^2+^ signals

The cholinergic anthelmintic levamisole is used to treat nematode parasite infections. We have previously demonstrated that the subunits for the levamisole sensitive nicotinic acetylcholine receptor (nAChR) are expressed in the intestine of *Ascaris suum* and that application of levamisole generates a distinctive Ca^2+^ signal that is inhibited using mecamylamine [21]. Hu et al. 2010b have observed that combination of levamisole and Cry5B has a synergistic killing effect on whole *C. elegans*.

We sought to determine if this effect could be detected as an interaction between Cry5B and levamisole on the *Ascaris suum* intestine using the approach that we have described above. We used 10 µg/ml Cry5B and 2-hour incubations and observed Ca^2+^ responses with amplitudes of 13.11% ± 0.51% like the 6-hour experiments with peaks occurring at an average time of 76.6 mins ± 2.7 mins, Figs. 8A, D &E; white bars. We observed with levamisole however, a reduction in the average % of the regions of the intestines responding to Cry5B after 2 hours compared to 6 hours, down to 76.4% ± 9.1%, Fig. 8F: white bar, suggesting that shorter term incubations result in fewer regions of the intestine responding.

**Figure 8:**
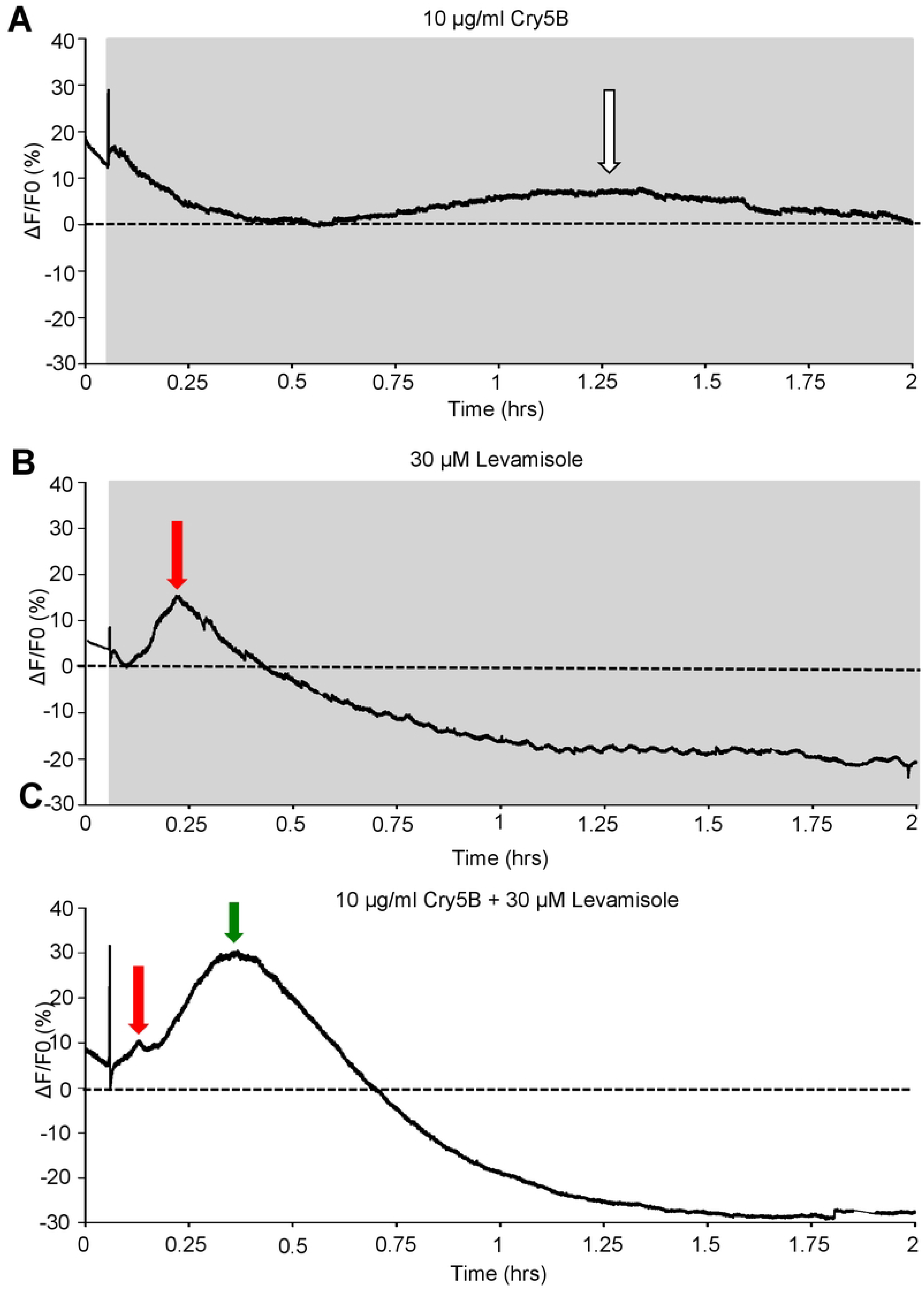
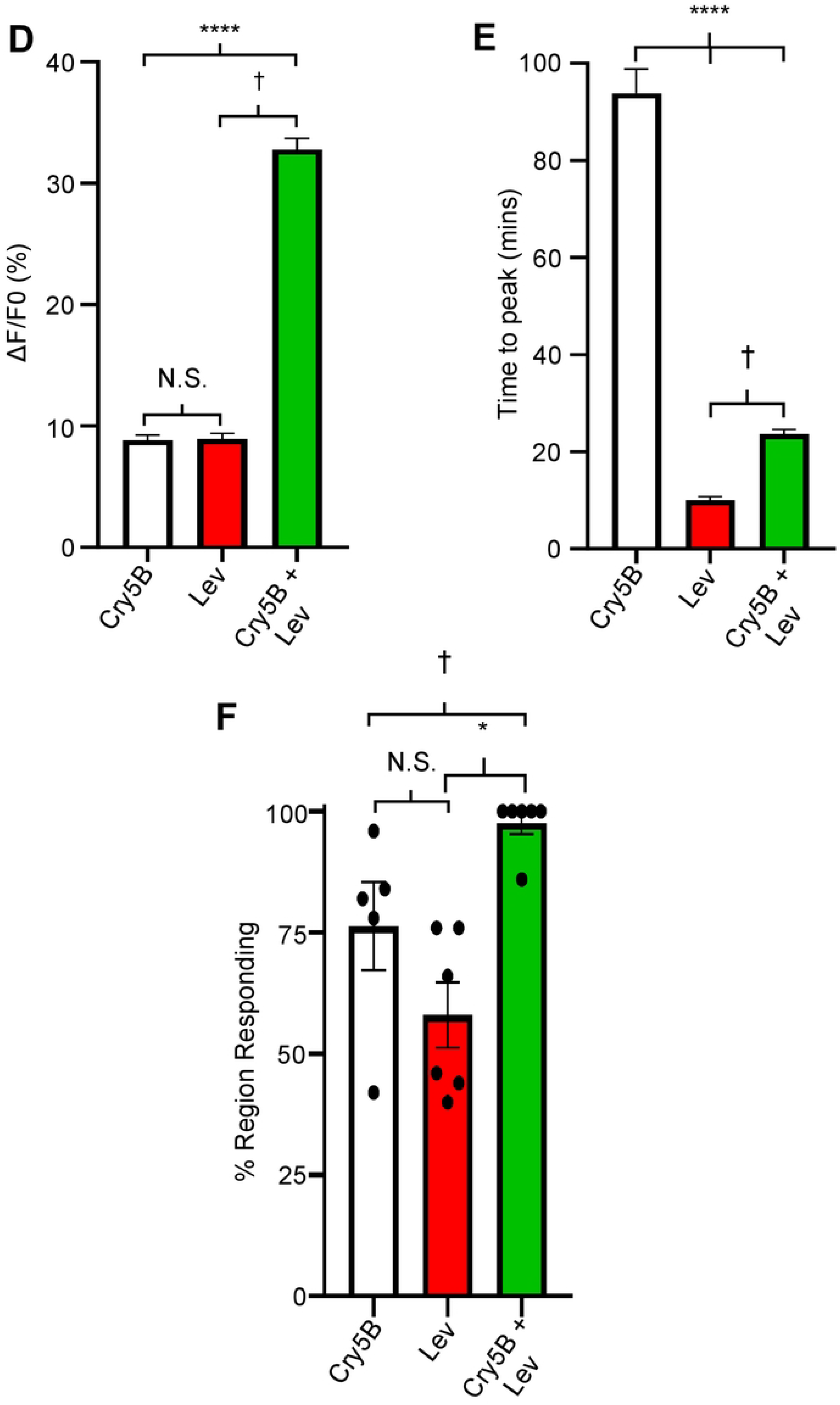
Levamisole potentiates Cry5B signal in *Ascaris* intestines: A) Representative trace highlighting a 2-hour recording in response to 10 µg/ml Cry5B. Grey box indicates Cry5B application and dotted line represents 0% F/F_0_. White arrow indicates peak of the Ca^2+^ signal. B) Representative trace highlighting a 2-hour recording in response to 30 µM levamisole. Grey box indicates levamisole application and dotted line represents 0% F/F_0_. The red arrow indicates the peak of the Ca^2+^ signal. C) Representative trace highlighting a 2-hour recording in response to the combination of 10 µg/ml Cry5B and 30 µM levamisole. The grey box indicates Cry5B, and levamisole application and dotted line represents 0% F/F_0_. Red arrow highlights the initial levamisole response and green arrow indicates the larger secondary response D) Average maximal amplitudes of Ca^2+^ fluorescence over 2 hours in 10 µg/ml Cry5B treated (white bar), 30 µM levamisole treated (red bar) and 10 µg/ml Cry5B + 30 µM levamisole treated (green bar). N.S. not significantly different to 10 µg/ml Cry5B (10 µg/ml Cry5B vs 30 µM levamisole, *P* = < 0.3117 *t* = 1.013, *df* = 393, unpaired *t*-test). **** Significantly different to 10 µg/ml Cry5B (10 µg/ml Cry5B vs 10 µg/ml Cry5B + 30 µM levamisole, *P* = < 0.0001 *t* = 13.58, *df* = 482, unpaired *t*-test). † Significantly different to 30 µM levamisole (30 µM levamisole vs 10 µg/ml Cry5B + 30 µM levamisole *P* = < 0.0001 *t* = 13.41, *df* = 465, unpaired *t*-test). E) Average time to maximal peak amplitude over 2 hours in 10 µg/ml Cry5B treated (white bar), 30 µM levamisole (red bar) and 10 µg/ml Cry5B + 30 µM levamisole (green bar). **** Significantly different to 10 µg/ml Cry5B (10 µg/ml Cry5B vs 30 µM levamisole, *P* = < 0.0001 *t* = 22.11, *df* = 363, unpaired *t*-test; 10 µg/ml Cry5B vs 10 µg/ml Cry5B + 30 µM levamisole, *P* = < 0.0001 *t* = 23, *df* = 482 unpaired *t*-test). † Significantly different to 30 µM levamisole (30 µM levamisole vs 10 µg/ml Cry5B + 30 µM levamisole *P* = < 0.0001 *t* = 12.37, *df* = 465, unpaired *t*-test). F) Average percent of regions showing a Ca^2+^ response larger than the average spontaneous Ca^2+^ amplitude in untreated intestines in 10 µg/ml Cry5B treated (white bar), 30 µM levamisole (red bar) and 10 µg/ml Cry5B + 30 µM levamisole (green bar). N.S. not significant compared to 10 µg/ml Cry5B (10 µg/ml Cry5B vs 30 µM levamisole, *P* = < 0.102 *t* = 1.821, *df* = 9, unpaired *t*-test). * Significantly different to 10 µg/ml Cry5B (10 µg/ml Cry5B vs 10 µg/ml Cry5B + 30 µM levamisole *P* = < 0.0357 *t* = 2.468, *df* = 9, unpaired *t*-test). † Significantly different to 30 µM levamisole (30 µM levamisole vs 10 µg/ml Cry5B + 30 µM levamisole, *P* = < 0.0004 *t* = 5.266, *df* = 10, unpaired *t*-test). 10 µg/ml Cry5B *n* = 5 individual female intestines with a total of 191/250 regions providing viable responses which were used to generate the mean and SEM for D & E. Note: 59/250 regions showed no response. 30 µM levamisole *n* = 6 individual female intestines with a total of 174/300 regions providing viable responses which were used to generate the mean and SEM for D & E. Note: 126/300 regions showed no response. 10 µg/ml Cry5B + 30 µM levamisole *n* = 6 individual female worms with a total of 293/300 regions providing a viable response were used to generate the mean and SEM for D & E. Note: 7/300 regions showed no response.

Application of 30 µM levamisole to the *Ascaris suum* intestine produces an early, ∼10min sharp peak, with an average amplitude around 10% [21]. We incubated intestinal tissues over 2 hours and recorded the levamisole signal to look for any longer-term effects of 30 µM levamisole exposure. The 2-hour levamisole Ca^2+^ responses were like the shorter levamisole applications. The peak Ca^2+^ response had an average amplitude of 12.4% ± 0.5% and time to peak of 11.5 mins ± 0.8 mins, Figs. 8B, 8D: red bar & 8E: red bar. After the initial peak, we saw no further major peaks other than small oscillations in the signal for the rest of the recording, Fig. 8A. These observations show that levamisole causes a more rapid influx of Ca^2+^ into the intestinal enterocytes. The levamisole mediated Ca^2+^ signal distribution was limited to 58% ± 7% of the intestine, Fig. 8F: red bar, suggesting that the levamisole receptors are not distributed evenly over the intestine surface.

With both 10 µg/ml Cry5B and 30 µM levamisole producing similar peak amplitudes, ∼12%, but having different rise times (11 mins vs 77 mins) we sought to determine if co-application of the two compounds altered the Ca^2+^ signal and/or affected the number of regions responding. Significantly, the combination of the two compounds resulted in a larger Ca^2+^ signal that had an average peak amplitude of 37.9% ± 1.4%, Fig. 8C & 8D: green bar, which took a significantly shorter time to reach its peak, 23.8 mins ± 0.6 mins, Fig. 8E: green bar compared to Cry5B alone. Although slower than the levamisole alone, this larger signal reveals the effect of their combination. Additionally, the combination of both 10 µg/ml Cry5B and levamisole resulted in 97.7% ± 2.3% of the regions tested showing a response, Fig. 8F: green bar, showing that the combination affects the intestine across more of the areas of the intestine than either of the compounds used alone. Our results highlight that the combination of both Cry5B and levamisole causes increased amplitudes and larger areas of activation compared to use of the either compound alone or faster rise to peak compared to Cry5B alone.

### Histological effects of the levamisole and Cry5B combination

As before, we treated intestines attached to our *Ascaris* body flaps over 2 hours in either 30 µM levamisole or 30 µM levamisole + 10 µg/ml Cry5B, and prepared histological slides to observe structural effects. The 2-hour 30 µM levamisole treated sample did show a level of change in the enterocytes. Fig. 7B; left panel, shows that treatment with levamisole caused vacuolation and some disruption throughout the intestine, but the vacuoles were more distributed and more apparent at the apical region of the cells. There was separation between some of the neighboring cells at the basolateral region of the preparation. We also observed evidence of damage to the brush border of the intestine, with small breaks appearing along the apical region of the enterocytes. These observations suggest that levamisole on its own can disrupt the integrity of the intestine with some unique phenotypes, illustrating a mode of action for this compound other than on the neuromuscular system of the nematode parasites.

Although levamisole and Cry5B alone caused some degree of intestinal damage, combination of the two compounds resulted in more severe destruction. As shown in Fig. 7B; right panel, enterocytes of intestines that were treated with both compounds showed more severe damage, with a more vacuoles, more tearing and loss of integrity throughout the intestine and increased evidence of necrotic cells. The increased histological damage is correlated with the bigger and faster Ca^2+^ signal suggesting that the enterocytes are showing necrotic cell-death at a faster rate than when either compound is used alone. The combination of Cry5B and the cholinergic anthelmintic levamisole leads to faster destruction of the *Ascaris* intestine that will contribute to a faster killing of the nematode parasites [13]

## Discussion

### Cry5B, Ca^2+^ and Necrosis

We observed message for *bre-5* [15, 16] required for the synthesis of arthroseries glycolipids, and *cdh-8* message for cadherin [17] present in the *Ascaris suum* intestine. Both the arthroseries glycolipids and CDH-8 have been described as Cry5B receptors. Once Cry5B has bound to these receptors, small 1-2 nm pores are formed in the plasma membrane which based on analogy to Cry1Aa and Cry1Ac pores are permeable to Ca^2+^ [29]. We found that Cry5B produced a slow, but large, dose-dependent increase in the Ca^2+^ signal that peaked at an average of 80 minutes across 95% of the intestine. The effects of Cry5B were inhibited by 100mM galactose which inhibits Cry5B binding to the glycolipids [15, 16] suggesting these are the major receptor targets in *Ascaris suum* intestines.

Histology showed significant damage produced by Cry5B at 2 hours and severe necrosis by 6 hours. The damage included vacuolation in the enterocytes, damage to the enterocyte brush borders and separations between adjacent enterocytes. These effects may be explained by a Ca^2+^ mediated necrosis cell-death pathway, Fig.9, that follows Ca^2+^ overload due to Ca^2+^ entry through Cry5B toxin pores formed in the plasma membranes of the enterocytes [17, 30].

**Figure 9:**
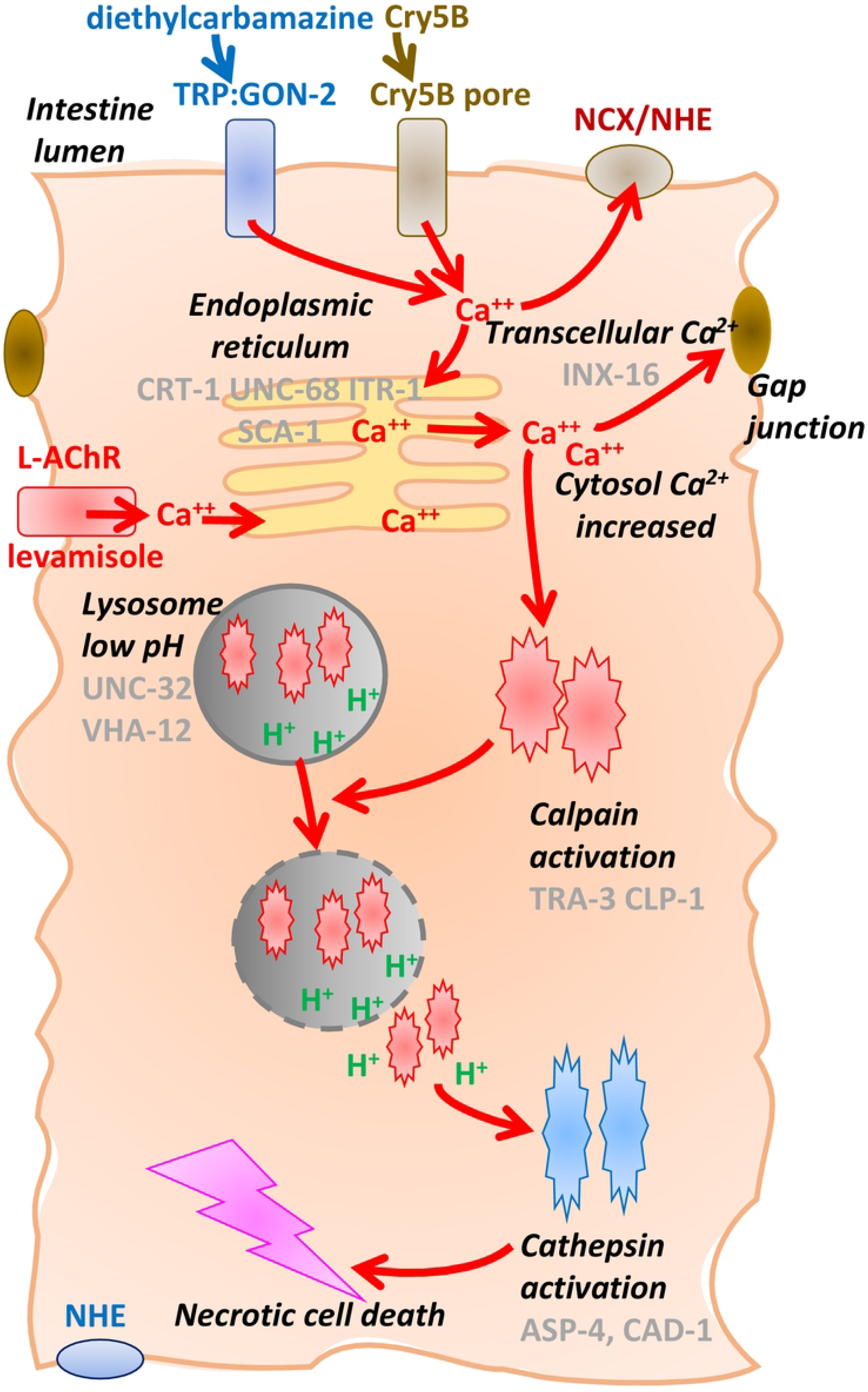
Summary diagram of proposed mode of action Cry5B and levamisole action on the intestine of *Ascaris suum* precipitating necrotic cell-death. High, uncontrolled increases in cytosolic Ca^2+^ in enterocytes leads to the necrosis cell-death pathway [30]. The increased Ca^2+^ can arise through Cry5B pores in the plasma membrane, opening of L-AChR channels by levamisole, activation of GON-2 TRP diethylcarbamazine activated channels or other unspecified pathways. The increase in cytosolic Ca^2+^ can pass through Gap junctions (INX-16) to adjacent enterocytes. Enterocyte cytosolic Ca^2+^ is controlled and affected by: 1) uptake by plasma and sarcoplasmic reticulum ATPases (SCA-1 and MCA-3) and the Ca^2+^ exchangers (NCX-1 and NHE); 2) release from ryanodine receptors (UNC-68), the IP3 receptors (ITR-1), and Ca^2+^ binding proteins (CRT-1); and 3) entry from the extracellular space through the plasma membrane. The rise in Ca^2+^ produces rupture of the lysosomes, release of cathepsins and calpains and then the necrosis cascade and cell-death pathway [33, 34]. The increase in Ca^2+^ activates the calpains TRA-3 and CLP-1 that give rise to cytosolic acidosis and lysosomal lysis. Lysosomes have a low internal pH due to two proton pumps, VHA-12 and UNC-32, and the rupture of lysosomes releases protons that activate the cathepsins, ASP-4 and CAD-1 driving the necrosis cascade. Once the cytosolic Ca^2+^ exceeds a sustained critical level, the necrosis cell-death pathway becomes irreversible and leads to destruction of the intestine and then the nematode.

### Levamisole

Levamisole is an agonist of nematode Ca^2+^ permeable L-AChR ion-channels [31] which are expressed in the *Ascaris suum* intestines [21] as well as muscle [32]. Levamisole produces a distinctive sharp Ca^2+^ peak in the enterocytes, at an average of 12 minutes after application to the intestine [21] but which is limited to 58% of the area of the intestine. This peak at 12 minutes is much slower than that which follows levamisole application to body muscle: it may therefore be due to a secondary Ca^2+^ release that follows opening of enterocyte L-type AChRs [21]. Nonetheless, the application of 10 µg/mg Cry5B and levamisole together shortened the onset time to the Cry5B peak and resulted in a larger Ca^2+^ peak signal that was seen at an average of 24 minutes across nearly all the area of the intestines. The Cry5B peak was slower than the levamisole peak but was much bigger than the 10 µg/ml Cry5B alone. The potentiating effects of levamisole on the effects of Cry5B are underpinned by these two agents acting on two separate Ca^2+^ permeable channels in the plasma membranes, both of which increase the calcium concentration in the enterocytes to produce cytotoxicity.

### Necrosis Cell-Death Cascade

In *C. elegans* an excessive and uncontrolled increase in cytosolic Ca^2+^ in enterocytes [30] leads to the necrosis cell-death pathway. The rise in Ca^2+^ produces rupture of the lysosomes, release of cathepsins and calpains and then the necrosis cascade and cell-death pathway Fig. 9 [33–35]. Fig. 9 shows the enterocyte cytosolic Ca^2+^ to be affected by: 1) uptake by plasma and sarcoplasmic reticulum ATPases (SCA-1 and MCA-3) and the Ca^2+^ exchangers (NCX-1 and NHE); 2) release from ryanodine receptors (UNC-68), release from IP3 receptors (ITR-1), and release from Ca^2+^ binding proteins (CRT-1); and 3) entry through the plasma membrane through diethylcarbamazine-sensitive TRP channels [22], levamisole activated L-AChR channels [21] and bacterial pores like Cry5B. Once the Ca^2+^ has increased it can spread to adjacent enterocytes through gap junctions (INX-16). The increase in Ca^2+^ activates the calpains TRA-3 and CLP-1 that give rise to cytosolic acidosis and lysosomal lysis. Lysosomes have a low internal pH due to two proton pumps, VHA-12 and UNC-32, and release of the protons activates the cathepsins, ASP-4 and CAD-1 driving the necrosis cascade. Once the cytosolic Ca^2+^ exceeds a sustained critical level the necrosis cell-death pathway becomes irreversible and leads to destruction of the intestine and then the nematode.

### Defense pathways against Cry5B

If the Ca^2+^ does not exceed the sustained critical level, the enterocytes can respond against this attack by activating defense pathways. Low concentrations of Cry5B, although they open Ca^2+^ permeable cation pores in cell plasma membranes, they may not kill *C. elegans* or nematode parasites. Nematodes can resist the effects of Cry5B by activating intrinsic cellular defense (INCED) pathways that include four components [36–40].

1. *A mitogen-activated p38/MAPKK (sek-1) pathway* of protein kinases, which is conserved in vertebrates. The pathway controls phosphorylation and activation of transcription factors driving the unfolded protein response to defend *C. elegans* against pore forming bacteria. The unfolded protein response (UPR) has a protective effect against Cry5B [39].
2. *A JNK MAPK pathway,* which is a ‘*key regulator*’ of the defense against pore-forming toxins that involves activation of the defense protein, AP-1 [36].
3. *A Rab-5/Rab-11 Vesicle-Trafficking pathway,* which is involved in endocytosis and membrane shedding of the plasma membrane [37]. Membrane trafficking transports proteins within the Golgi body to the plasma membrane and requires RAB-5 and RAB-11 proteins. These proteins are required for the defense against pore-forming toxins like Cry5B and repair of plasma membranes.
4. *An actin pathway,* which is also involved in the defense of epithelial cells like the enterocytes of the intestine. Actin cytoskeletons within the enterocyte are present in a dense network under the microvilli of the apical surface. Pore-forming toxins interact with the actin filaments to promote uptake of the protoxin by endocytosis. These pore-forming toxins also act directly on actin, to produce migration and/or breakdown of actin from the apical surface of the enterocytes to the basolateral surface which results in gaps between adjacent cells and increased permeability of the intestine. [41–43]. The actin related gene *nck-1* functions with F-actin cytoskeleton modifying genes like *Arp2/3*, *erm-1, dbn-1* and *nck-1/arp2/3* to promote pore repair of the membrane and is important for Cry5B defense [44]. Loss of nck-1 gives rise to Cry5B hypersensitivity.

### Defense pathways against levamisole

Less is known about the effects of maintained concentrations of levamisole on nematode intestines but studies on the effects of maintained concentrations of levamisole on nematode muscle have been observed in *Brugia malayi* [45] and *C. elegans* [46]. The effects of maintained concentrations of levamisole following spastic paralysis leads to flaccid paralysis and then a recovery of motility through processes that allows for homeostatic plasticity (accommodation and adjustment to maintained stimuli). In *B. malayi* there is a loss of functional L-AChR channels an increase in levamisole resistant AChRs, and a decrease in *nra-2* (a nicalin homologue that encodes an ER retention protein) that allows faster loss of sensitivity to levamisole and recovery. The regulators of desensitization and therefore defense and recovery of AChRs that are present in *C. elegans* muscle and *Brugia* include: TAX-6, a calcineurin A subunit, that affect desensitization of AChRs in rat chromaffin cells; SOC-1 (a multi-subunit adaptor protein); and PLK-2 (a serine/threonine kinase) [46–48].

### How do the levamisole and Cry5B interact?

In *C. elegans* there is a synergistic interaction between levamisole and Cry5B that displays hyper-susceptibility [18]. We have seen that both individual agents can lead to Ca^2+^ cytotoxicity of the *Ascaris* enterocytes. If one of these compounds induces a cytoplasmic Ca^2+^ near a critical level, then the addition of the other agent will trigger the necrosis cell death pathway. If the cytoplasmic Ca^2+^ are not sufficient to trigger the necrosis cell-death pathway, the defense pathways will permit the enterocytes to recover and overcome the effects of the Cry5B and levamisole.

### Hyper-susceptibility

Homeostatic plasticity, a general physiological phenomenon that is found in both vertebrates and invertebrates [45, 49] whereby tissues adjust their excitability and responses (providing they do not undergo cell death) to maintained concentrations of Ca^2+^ when they are either too low or two high for low. The plasticity involves changes in the number and distribution of membrane ligand-gated ion-channels, voltage-activated ion-channels, calmodulin, calcineurin, intracellular calcium release and uptake [49–51]. Thus, a low intracellular Ca^2+^ concentrations produced with levamisole receptor mutants is anticipated to make the tissues more sensitive to Cry5B than WT *C. elegans*.

## Conclusion

We have observed that levamisole and Cry5B produce an increase in Ca^2+^ in the enterocytes of the *Ascaris suum* intestine and lead to necrosis and cell-death. The effects of the two compounds are synergistic, being much bigger and sooner in onset with both are administered together. A combination of the two compounds is expected to be more effective than either of the two compounds alone and be more effective in the presence of resistance to one of these compounds.

## Acknowledgements

We acknowledge the NIH National Institute of Allergy and Infectious Diseases grants R01AI047194 and R01AI155413 to RJM, the E. A. Benbrook Foundation for Pathology and Parasitology for support to RJM. The funding agencies had no role in the design, execution or publication of this study. The content is solely the responsibility of the authors and does not necessarily represent the official views of the National Institute of Allergy and Infectious Diseases. We acknowledge Yan Hu and Kelly Flanagan for the purification and preparation of the Cry5B. We finally acknowledge Tracey Stewart in generating and reviewing our histology slides.

## Author contributions

P. W. and R. M. wrote the main manuscript text. P.W., R.M., and A.P.R. provided interpretations, conceptions, and designs of experimental procedures. P.W. acquired and analyzed experimental data. M.B. reviewed histology slides. All authors have reviewed the manuscript.

## Competing interests

The authors declare no competing interests.

---

**Supplementary Figure 1.**
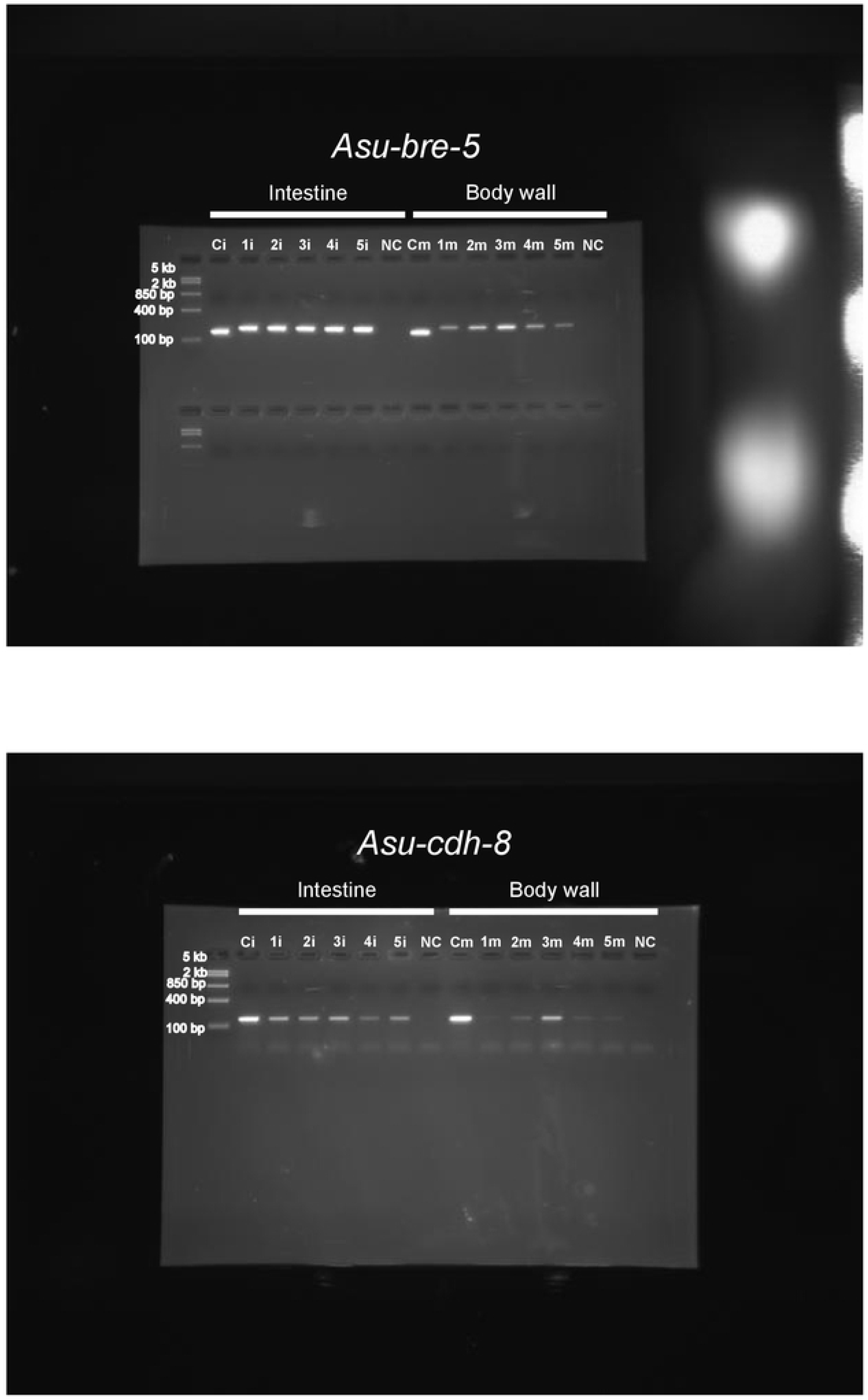

**Supplementary Table 1.**
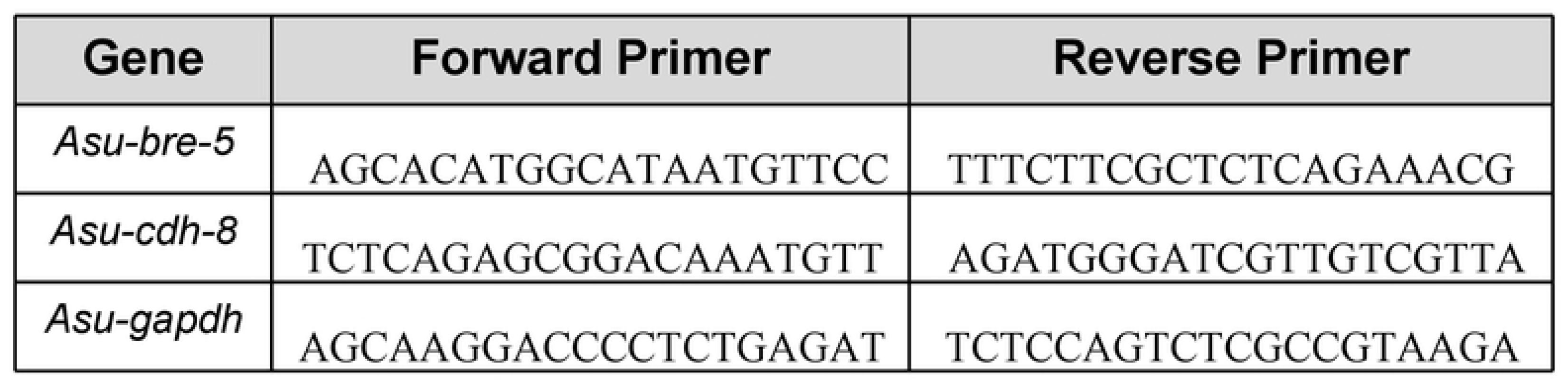

**Supplementary Table 2.**
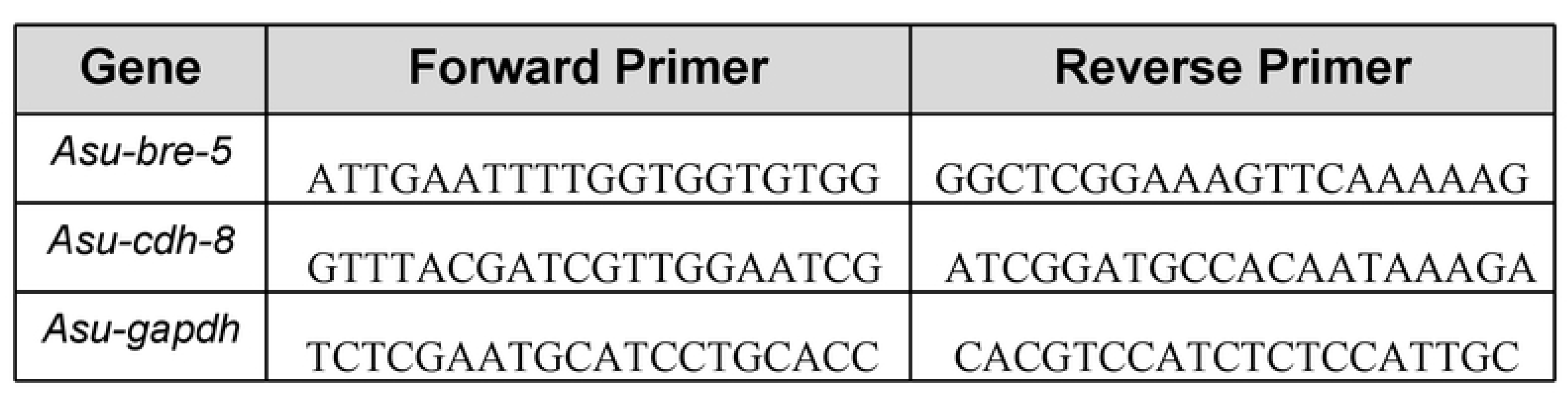

**Supplementary Table 3.**

| Gene | Accession number |
| --- | --- |
| *Asu-bre-5* | AgB01_g494/AEUI03000001.1 |
| *Asu-cdh-8* | AgR004_g364/AEUI03000009.1 |

## Notes

### Competing Interest Statement

The authors have declared no competing interest.

